# Generation of Cellular Biofactories for Scalable Production of Surface-Engineered Extracellular Vesicles via CRISPR Genome Editing

**DOI:** 10.1101/2025.09.30.679598

**Authors:** Yuki Kawai-Harada, Thomas Scarborough, Nayeema Siraj, Jayadeep Yedla, Tiffany Rennells, S. Patrick Walton, Christina Chan, Masako Harada

**Affiliations:** Institute for Quantitative Health Science and Engineering (IQ), Division of Chemical Biology, Michigan State University; Department of Biomedical Engineering, Michigan State University; Department of Chemical Engineering and Material Science, Michigan State University; Department of Microbiology, Genetics, & Immunology, Michigan State University; Lyman Briggs College, Michigan State University; Department of Physiology, Michigan State University; Department of Biochemistry and Molecular Biology, Michigan State University

**Keywords:** Engineered extracellular vesicles (EVs), CRISPR-Cas9, Stable cell line, EV surface engineering, Scalable EV production

## Abstract

Extracellular vesicles (EVs) are versatile biological nanoparticles with applications in therapeutics, diagnostics, and biotechnology. Current production methods using transient transfection or chemical conjugation suffer from high variability, limited scalability, and heterogeneous EV populations. Here, we developed CRISPR-Cas9 engineered HEK293T cell lines with stable integration of mCherry-C1C2 fusion proteins at the AAVS1 locus for continuous production of surface-modified EVs. The engineered cell lines demonstrated significantly higher surface display efficiency compared to transient transfection, with reduced batch-to-batch variability. EVs maintained native characteristics including size distribution (120-130 nm) and marker expression while showing efficient cellular uptake. The platform maintained consistent production of uniformly modified EVs with stable transgene expression over at least 25 passages (~3 months), eliminating the need for repeated transfections and reducing batch-to-batch variability inherent to transient expression systems.

## Introduction

Extracellular vesicles (EVs) represent highly evolved delivery systems, yet their clinical translation remains hampered by fundamental manufacturing challenges. These membrane-bound nanoparticles, secreted by virtually all cell types, represent a conserved mechanism of intercellular communication^1^. Their intrinsic biocompatibility, low immunogenicity, and capacity to transport molecules between cells surpass synthetic alternatives, making EVs promising candidates for innovative therapeutic applications^2^. The EV family encompasses vesicles of diverse origins and biogenesis pathways. Exosomes (30-150 nm) originate from the endosomal pathway through multivesicular body (MVB) formation, while microvesicles (MVs: 100-1000 nm) bud directly from the plasma membrane^3,4^. Despite this heterogeneity, EVs share key features that make them attractive for controlled delivery applications: enrichment of targeting receptors, protection of cargo from degradation, and natural mechanisms for cellular uptake^5,6^. The surface display of phosphatidylserine (PS) on EVs provides both a recognition signal for recipient cells and a platform for engineering approaches^7^. Unlike phosphatidylcholine (PC), the most abundant membrane phospholipid, PS is selectively externalized on EV surfaces where it serves functional roles in cellular uptake^8,9^. PS-mediated interactions with receptors such as Tim4 facilitate EV internalization^9^, and surface-exposed PS has been shown to enhance uptake by specific cell types, including liver macrophages^8^. This selective PS externalization makes PS-binding domains particularly attractive for engineering EVs with modulated cellular uptake and targeting properties.

The therapeutic potential of EVs has attracted significant clinical interest, and early clinical successes — including MSC-derived EVs for inflammatory diseases — have validated the therapeutic concept^10^. However, the transition from laboratory proof-of-concept to scalable manufacturing has revealed significant challenges for clinical translation. Genetic engineering and chemical modifications are the most widely used methods for EV engineering. These approaches enable the modification of EV surfaces for specific targeting and the loading of therapeutic cargo for enhanced functionality^11^. Both approaches, while useful for mechanistic studies, produce heterogenous EV populations with variable modification efficiency. Genetic manipulation, typically achieved through transient transfection, introduces gene cassettes that fuse targeting peptides to membrane-associated proteins such as LAMP2B^12-16^, tetraspanins (CD9, CD63, CD81)^17-19^, or to phosphatidylserine-binding bridging proteins like lactadherin (MFGE8)^20-22^. This approach enables the surface display of targeting ligands and remains one of the most widely adopted strategies for engineering EVs. However, it faces key challenges for therapeutic applications, including heterogeneous modification across EV populations and the transient nature of engineered EV (eEVs) production^20-22^. Indeed, some studies report that only ~50% of EVs produced through transfection display the intended modification^23^, underscoring the limitation of this method. Chemical modification offers an alternative, enabling direct conjugation of ligands onto the EV surface. Covalent methods such as click chemistry exploit the reaction between alkyne-functionalized EVs and azide-functionalized ligands to achieve stable linkages^24^. Non-covalent approaches, such as the insertion of lipid-anchored ligands (e.g., DSPE-PEG), allow spontaneous incorporation into EV membranes while presenting targeting moieties on the surface^25^. Although chemical strategies often achieve more consistent modifications than genetic methods, they can alter EV biophysical properties^25^ and remain limited in scalability and long-term production. To overcome these challenges, we investigated the C1C2 domain of lactadherin, which binds PS with high affinity and has demonstrated efficient surface display in our studies^22,26^. Nevertheless, like most approaches in the field, these methods rely on transient expression systems that compromise manufacturing consistency.

Controlled release research has long emphasized that successful therapeutic translation requires robust manufacturing processes that ensure product quality and consistency^27^. For EV-based therapeutics, this presents unique challenges. Unlike synthetic drug delivery systems, where manufacturing parameters can be precisely controlled, EV production depends on complex biological processes that are inherently variable^28^. Additionally, EV storage and handling present significant reproducibility challenges, as our previous work demonstrated that standard PBS storage compromises EV integrity and function^29^. Current good manufacturing practice (cGMP) guidelines for cell and gene therapies provide a framework for EV production, but existing methods fall short of these standards^30^. For example, transient transfection approaches require repeated interventions, introduce batch-to-batch variability, and scale poorly. Regulatory agencies increasingly emphasize the need for well-characterized, consistently produced materials^31^.

Stable cell line platforms offer a promising solution by enabling sustained, uniform expression of engineered components. Traditional approaches using random integration suffer from positional effects and epigenetic silencing that compromise long-term stability^32,33^. The development of CRISPR-Cas9 technology has transformed this landscape by enabling precise, targeted integration into genomic safe harbors^34^. The Adeno-Associated Virus Integration Site 1 (AAVS1) locus has emerged as the golden standard for stable transgene expression in human cells. Located within intron 1 of the PP1R12C gene, this site supports robust expression without disrupting endogenous function^35^. Multiple studies have demonstrated stable expression over 30 passages or 3-6 months while maintaining transgene activity^12,36^. For EV manufacturing applications, AAVS1 targeting offers the potential to create cellular “biofactories” that produce consistently eEVs. Despite these advances, few studies have systematically evaluated stable cell lines for EV engineering applications.

This study therefore focuses on evaluating how stable expression influences EV production and surface modification efficiency, whether consistent quality can be maintained over extended culture periods necessary for therapeutic manufacturing, and the potential advantages of stable cell lines in modification density and functional performance compared to transient systems. We hypothesized that targeted integration of our mCherry-C1C2 fusion construct into the AAVS1 locus would provide a stable platform that outperforms transient methods in both modification efficiency and long-term consistency. The C1C2 domain’s high affinity for PS enables efficient EV surface display, while mCherry fusion allows real-time monitoring of both cellular expression and EV modification. By targeting the AAVS1 locus, we aim to minimize positional effects that can compromise stable expression, providing a foundation for consistent production. Beyond demonstrating feasibility, this study systematically evaluates these engineered cell lines from a manufacturing perspective, assessing scalability, modification efficiency, and functional consistency. Our results demonstrate that CRISPR-engineered stable cell lines achieve efficient surface modification sustained expression over extended culture periods, establishing a reproducible platform for manufacturing uniformly modified EVs.

## Materials and Methods

### DNA Construct

The AAVS1 T2 CRIPR in pX330 was a gift from Masato Kanemaki (Addgene plasmid # 72833; http://n2t.net/addgene:72833; RRID:Addgene_72833) and was used to target the AAVS1 locus as sgRNA-Cas9 plasmid^37^. Donor plasmid, pDNR-CMV-mCherry-C1C2, was constructed from AAVS1 EGFP-Sec61B that was a gift from Jens Schmidt (Addgene plasmid # 207553; http://n2t.net/addgene:207553; RRID:Addgene_207553) by replacing promoter and gene cassette with CMV-mCherry-C1C2 that was amplified from pcS-mCherry-C1C2 (Addgene #178425)^38^. pDNR-EF-1α-mCherry-C1C2 was constructed by amplifying the EF-1α promoter sequence from human genomic DNA by PCR and replacing it with the CMV promoter on pDNR-CMV-mCherry-C1C2.

### Cell Culture and Stable Cell Line Generation

Cell lines, HEK293T (Human Embryonic Kidney cell line) obtained from American Type Culture Collection (ATCC), EGFP-expressing H1299 (generously provided by Dr. Jørgen Kjems and Dr. Manfred Gossen^39^) were routinely tested for mycoplasma contamination. Cells were cultured in high-glucose DMEM (Gibco) supplemented with 100U/mL penicillin, 100 µg/mL streptomycin, and 10% (v/v) fetal bovine serum (FBS, Gibco), and maintained in a humidified incubator with 5% CO2 at 37°C. Nucleofection was used to introduce DNA into cells. 1 million cells were suspended in Nucleofection buffer (5mM KCl, 15mM MgCl2, 15mM HEPES, 150mM Na2HPO4/NaH2PO4 (pH7.2), 50mM Mannitol) ^40^ together with sgRNA-CRISPR plasmid and donor plasmid at 1:1 ratio (w/w), and 2 µg DNA was introduced into the cells using 4D-Nucleofector (Lonza) to construct a stable cell line pool. To obtain CRISPR-positive cells, selection was performed with 1 μg/mL of Puromycin for 2 weeks, followed by PCR with primer for junction (F:ACCAACGCCGACGGTATCAG/R:CAGACCCTTGCCCTGGTGGT) to confirm that the bulk of cells included CRISPR-positive cells. Single clones were obtained from the stable cell line pool by Fluorescence-Activated Cell Sorting (FACS) using the MA900 cell sorter (SONY) in the Michigan State University Flow Cytometry Core Facility, and gene insertion into the AAVS1 locus was confirmed by genotyping PCR with primer for junction and insertion site (F:CCTTACCTCTCTAGTCTGTGCTA/R:GAGAGATGGCTCCAGGAAATG).

### EV Isolation

The cells were seeded in high-glucose DMEM (Gibco) supplemented with 100 U/mL penicillin, 100 µg/mL streptomycin (Pen-Strep, Lonza), and 10% (v/v) fetal bovine serum (FBS, Gibco) one day before media replacement with DMEM supplemented with insulin-transferrin-selenium (ITS, Corning) and Pen-Strep. 48 hours after media replacement, the conditioned media were collected for EV isolation. As a transient transfection control, HEK293T cells were transfected with pcS-mCherry-C1C2 using PEI 24 h prior to media change as described previously^22^. EVs were purified from conditioned media by differential centrifugation. Briefly, culture media were centrifuged at 600 xg for 15 minutes to remove cells and cellular debris, and the supernatant was further centrifuged at 2000 xg for 30 minutes to remove apoptotic bodies. The supernatant was then ultracentrifuged in PET Thin-Walled ultracentrifuge tubes (Thermo Scientific 75000471) at 100,000 xg with a Sorvall WX+ Ultracentrifuge equipped with an AH-629 rotor (k factor = 242.0) for 90 minutes at 4°C to pellet the EVs. The pellet containing EVs was resuspended in PBS. EVs were also isolated by Tangential Flow Filtration (TFF) system, μPULSE (Formulatrix) with 300kDa mPES filter chip. EVs were isolated using the default concentration program, followed by buffer exchange into PBS using the manufacturer’s buffer exchange program.

### Western Blotting

Cells were lysed using MRIPA lysis buffer (150 mM NaCl, 1.0% Triton X-100, 0.25% sodium deoxycholate, 50 mM Tris, pH 7.4), and the supernatant was collected as cell lysates. Protein samples (40 µg) were denatured at 95°C for 10 minutes in 1x Sample Buffer (50 mM DTT, 0.0025% bromophenol blue, 62.5 mM Tris-HCl [pH 6.8], 10% glycerol, 2% SDS), separated by SDS-PAGE on 4– 20% Mini-PROTEAN® TGX™ Precast Protein Gels (Bio-Rad), and transferred to PVDF membranes using a CAPS-based transfer buffer. EVs (2×10^9^ particles) were denatured at 70°C for 10 minutes in 1x NuPAGE LDS Sample Buffer (Thermo Fisher Scientific), separated by SDS-PAGE on 4–20% Mini-PROTEAN® TGX™ Precast Protein Gels (Bio-Rad), and transferred to PVDF membranes using a CAPS-based transfer buffer.

Membranes were blocked with 5% dry milk in TBS containing 0.1% Tween 20 (TBST) for 2 hours and then incubated with primary antibodies at 4°C overnight. Following three washes with TBS containing TBST, membranes were incubated with horseradish peroxidase-conjugated secondary antibodies for 2 hours at room temperature. After three additional TBST washes, protein bands were visualized using SuperSignal West Pico PLUS chemiluminescent substrate (Thermo Scientific) and imaged with ChemiDoc Imaging System (Bio-Rad). Primary antibodies used: anti-HA (Sigma Aldrich, H3663, RRID:AB_262051), anti-CD63 (Thermo Fisher, 10628D, RRID:AB_2532983), anti-ALIX (Proteintech, 12422-1-AP, RRID:AB_2162467), anti-Calnexin (Abcam, ab133615, RRID:AB_2864299), and anti-GAPDH (Cell Signaling Technology, #2118, RRID:AB_561053). Secondary antibodies used: Goat anti-Mouse IgG (H+L) Highly Cross-Adsorbed Secondary Antibody, HRP (Invitrogen, A16078, RRID:AB_2534751) and Goat anti-Rabbit IgG (H+L) Highly Cross-Adsorbed Secondary Antibody, HRP (Cell Signaling Technology, A16110, RRID:AB_2534783).

### Nanoparticle Tracking Analysis

The particle size and concentration were measured using a ZetaView® (Particle Metrix) Nanoparticle Tracking Analyzer following the manufacturer’s instructions. The following parameters were used for measurement: [Post Acquisition parameters (Min brightness: 22, Max area: 800, Min area: 10, Trace length: 12, nm/Class: 30, and Classes/Decade: 64) and Camera control (Sensitivity: 85, Shutter: 250, and Frame Rate: 30)]. EVs were diluted in PBS between 20- and 200-fold to obtain a concentration within the recommended measurement range (0.5 × 10^5^ to 10^10^ cm^−3^).

### Confocal Microscopy

5 × 10^4^ each of H1299 cells were seeded in 8-well chamber slide (0030742036, Eppendorf, Germany) 24 hours before EV treatment. The cells were incubated with 2 × 10^8^ previously isolated eEVs for 4 hours, followed by three PBS washes to remove unbound EVs. Cells were fixed with 4% PFA at room temperature for 15 minutes, washed with PBS containing 0.1% Tween 20 three times, and cells were incubated with DAPI for 10 minutes at room temperature. Slides were mounted with mounting medium (00-4958-02 Fisher Scientific, USA) and fluorescence images were taken at 60× magnification using confocal laser scanning microscopy (A1 HD25/A1R HD25 confocal microscope, Nikon, Japan) in the Center for Advanced Microscopy, Michigan State University.

### Immuno-TEM

Formvar/carbon film-coated 200 mesh gold EM grids (Electron Microscopy Sciences, FCF-200-AU) were soaked in 50 µL EVs for 30 min to allow adsorption onto the grid. The grids were then fixed with 50 µL of 2% paraformaldehyde (PFA) for 5 min and rinsed three times with 50 µL TEM-grade PBS. To quench free aldehyde groups, the grids were treated with 50 µL of 50 mM glycine for 10 min. Blocking was performed on 50 µL PBS containing 1% BSA for 30 min. After blocking, the grids were incubated with 50 µL of either anti-HA (Sigma-Aldrich, H3663) or anti-CD63 (Thermo Fisher, 10628D) antibody (1:100 in PBS with 0.1% BSA) for 1 h. The grids were then washed five times with 50 µL PBS containing 0.1% BSA, 10 min each. For secondary antibody labeling, the grids were incubated with 50 µL Goat-anti-Mouse IgG conjugated with 10 nm gold nanoparticles (Electron Microscopy Sciences, 25129), diluted 1:100 in PBS with 0.1% BSA, for 1 h. This was followed by three 10-min washes on 50 µL PBS containing 0.1% BSA and two washes with 50 µL HPLC-grade water. EVs were negatively stained with 25 µL of 1% uranyl acetate for 5 minutes. Grids were air-dried for 24–48 h, and images were acquired using a JEOL 1400 TEM at 100 kV.

## Results

### CRISPR-Cas9 Strategy for Stable Cell Line Generation

To establish stable cellular platforms for eEV production, we designed donor plasmids targeting the AAVS1 safe harbor locus (Fig. 1A). The donor construct contained left and right homology arms flanking an expression cassette with either CMV or EF-1α promoters driving mCherry-C1C2 fusion protein expression, followed by a puromycin resistance gene for selection. The AAVS1 locus was selected as the integration site due to its established properties as a genomic safe harbor that supports robust transgene expression without disrupting endogenous cellular functions^35^. Unique restriction enzyme sites were incorporated upstream of the promoter and flanking the gene cassette to facilitate future modifications.

**Figure 1.**
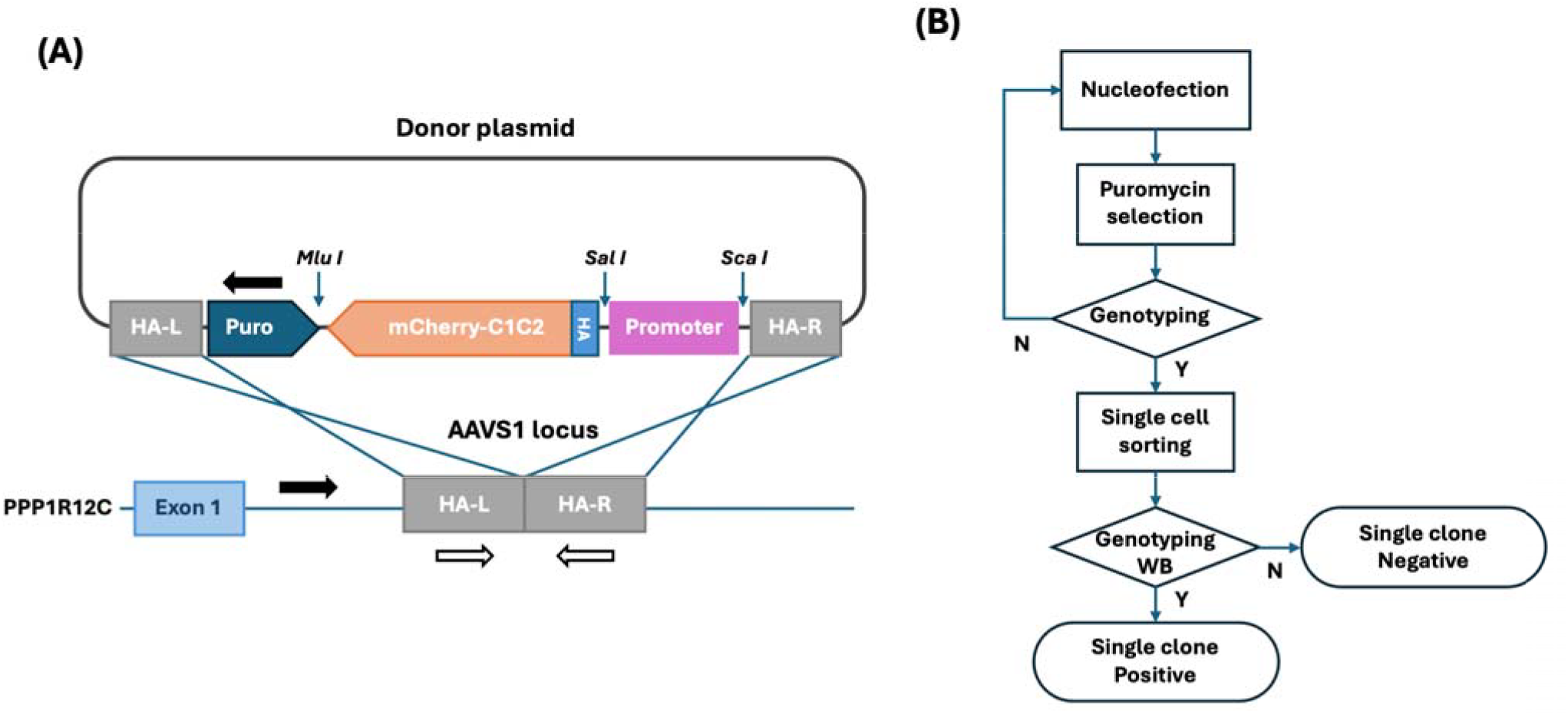
Design of donor plasmids for the AAVS1 locus and workflow for stable cell line generation. **(A)** Schematic diagram of gene expression construct targeting AAVS1 site. The donor plasmid contains left and right homology arms (HA-L and HA-R) flanking the expression cassette, which includes either CMV or EF-1α promoter driving mCherry-C1C2 fusion protein expression, followed by a puromycin resistance gene for selection. Black arrows indicate genotyping primers for junction PCR (confirming successful integration), while white arrows indicate primers for insertion site PCR (detecting wild-type allele). **(B)** Workflow for generating single clone stable cell lines via CRISPR-Cas9 genome editing. HEK293T cells are co-transfected with sgRNA-Cas9 and donor plasmids via nucleofection, followed by puromycin selection (1 μg/mL for 14 days) to enrich CRISPR-positive cells. Bulk cells are screened by junction PCR, and mCherry-positive single cells are isolated by fluorescence-activated cell sorting (FACS). Individual clones are validated by genotyping PCR and Western blot analysis for transgene expression.

The systematic workflow for generating stable cell lines is outlined in Figure 1B. HEK293T cells were co-transfected with sgRNA-Cas9 and donor plasmids using an optimized nucleofection buffer^40^. CRISPR-positive cells were subsequently enriched via puromycin selection (1µg/mL for 14 days). Bulk cell populations were screened by junction PCR to confirm successful integration before proceeding to single-cell isolation. mCherry-positive single cells were isolated by fluorescence-activated cell sorting (FACS), and individual clones were validated through genotyping PCR and western blot analysis.

### Successful Generation and Characterization of Single Cell Clones

Puromycin selection effectively eliminated untransfected cells, with complete cell death observed at concentrations ≥1 μg/mL after 14 days. Junction PCR analysis of bulk populations confirmed successful gene cassette insertion at the target site (Fig. S1). Single-cell clones were obtained by FACS sorting mCherry-positive cells into 96-well plates (Fig. S2). Genotyping analysis revealed distinct integration patterns between promoter systems (Fig. 2A). From CMV-mCherry-C1C2 transfected cells, 17 single clones were obtained, of which 7 showed successful gene insertion by genotyping PCR. All 7 positive clones were heterozygous for the integration event. In contrast, EF-1α-mCherry-C1C2 transfection yielded 45 single clones, with 35 showing successful integration (12 heterozygous and 23 homozyous integration clones). The genotyping strategy using junction PCR (confirming integration) and insertion site PCR (detecting wild-type alleles) effectively distinguished between heterozygous and homozyous integration events, consistent with established AAVS1 targeting protocols ^41^.

**Figure 2.**
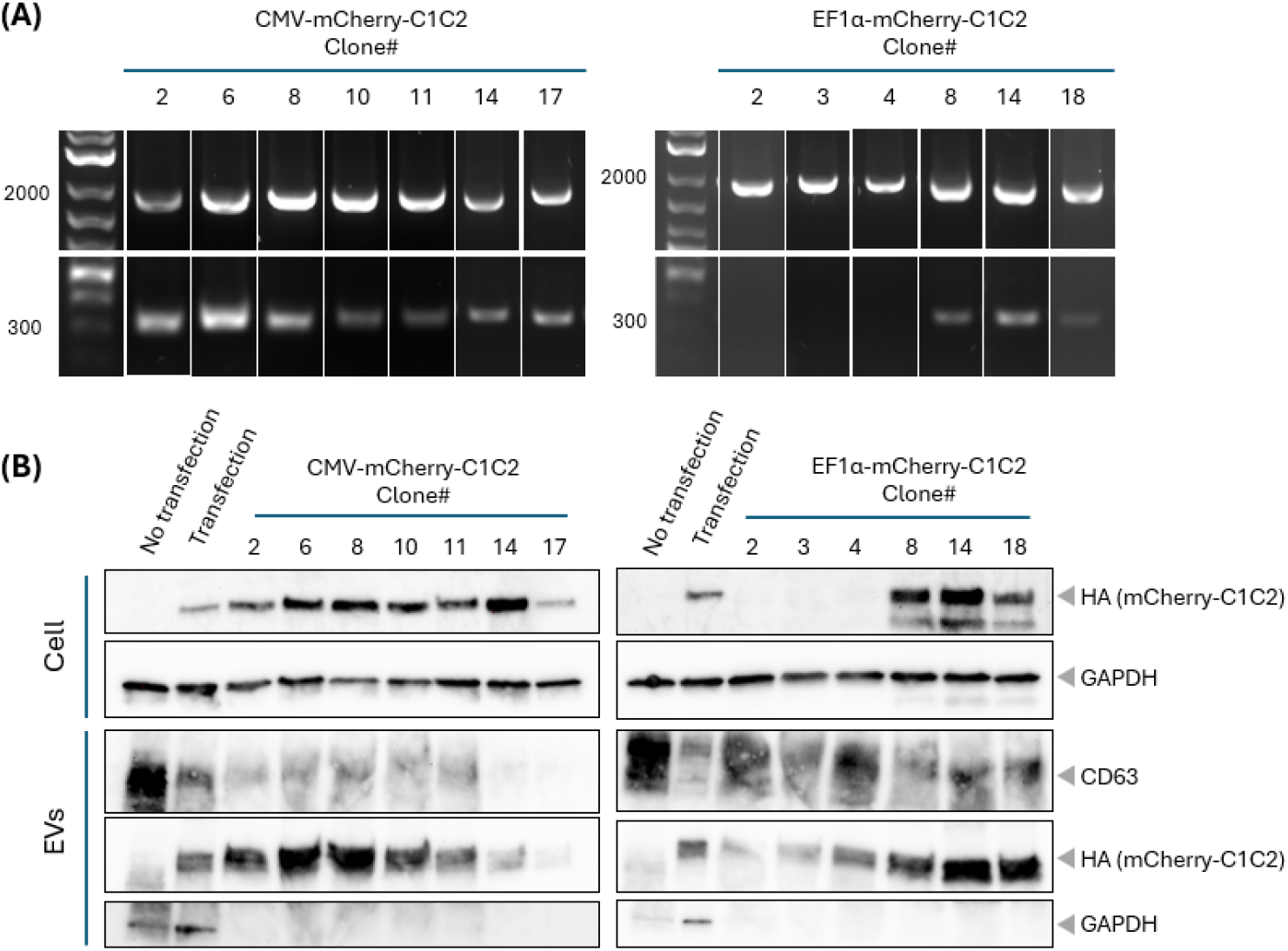
Genotyping of single clones and comparison of gene expression among them. **(A)** Representative genotyping PCR results for selected single clones. Top panel: Junction site PCR using primers spanning the integration site confirms successful CRISPR-mediated insertion (CRISPR-positive marker). Bottom panel: Insertion site PCR targeting the wild-type AAVS1 locus distinguishes heterozygous (both bands present) from homozygous integration clones (wild-type band absent). Numbers indicate clone identifiers for CMV-mCerry-C1C2 (clones 1-7) and EF-1α-mCherry-C1C2 (clones 8-13) cell lines. **(B)** Western blot analysis comparing mCherry-C1C2 expression levels in cell lysates (40 μg protein) and corresponding EVs (15 μg protein) from selected clones. Anti-HA antibody detects the HA-tagged mCherry-C1C2 fusion protein. GAPDH serves as loading control for cell lysates and as an endogenous PS-binding protein control for EVs. Clone #8 (CMV promoter) and clone #14 (EF-1α promoter) showed strong mCherry-C1C2 expression in EVs by Western blot bands and were selected for detailed characterization. Transient transfection control shows expression from pcS-mCherry-C1C2 plasmid 48h post-transfection.

Western blot analysis of selected clones revealed variable mCherry-C1C2 expression levels in both cellular and EV fractions (Fig. 2B). Expression levels in cells correlated with those in EVs across different clones. Notably, among EF-1α-driven clones, heterozygous clones showed higher expression levels than homozygous clones. Some EF-1α clones showed additional N-terminal fragments of the fusion protein in cellular lysates, though these did not affect EV modification (Fig. S3). Importantly, mCherry-C1C2 expression levels in stable cell lines were consistently higher than those achieved by transient transfection using the same construct^22^. Conversely, GAPDH levels in EVs showed an inverse correlation with mCherry-C1C2 expression, suggesting competition for PS-binding sites as previously described for PS-binding proteins^42^.

### Comprehensive Characterization of eEVs from Stable Cell Lines

Further detailed characterization was performed using CMV-mCherryC1C2 clone #8 and EF-1α-mCherryC1C2 clone #14 based on their higher expression levels in EVs. Fluorescence microscopy revealed that mCherry-C1C2 protein localized to punctate cytoplasmic structures consistent with endosomal and multivesicular body compartments involved in EV biogenesis (Fig. 3A), similar to subcellular distributions observed for other EV-targeted proteins^5^.

**Figure 3.**
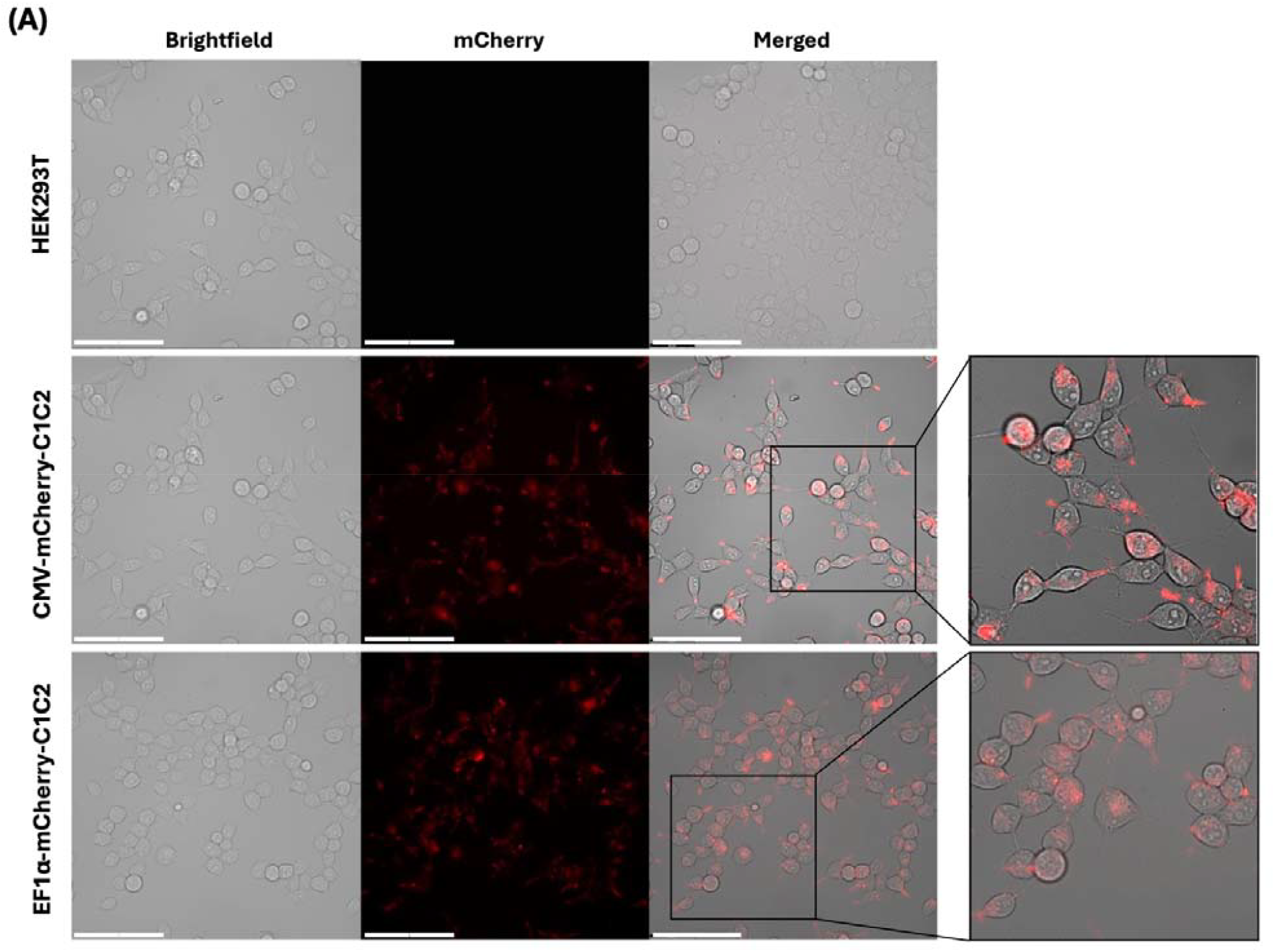
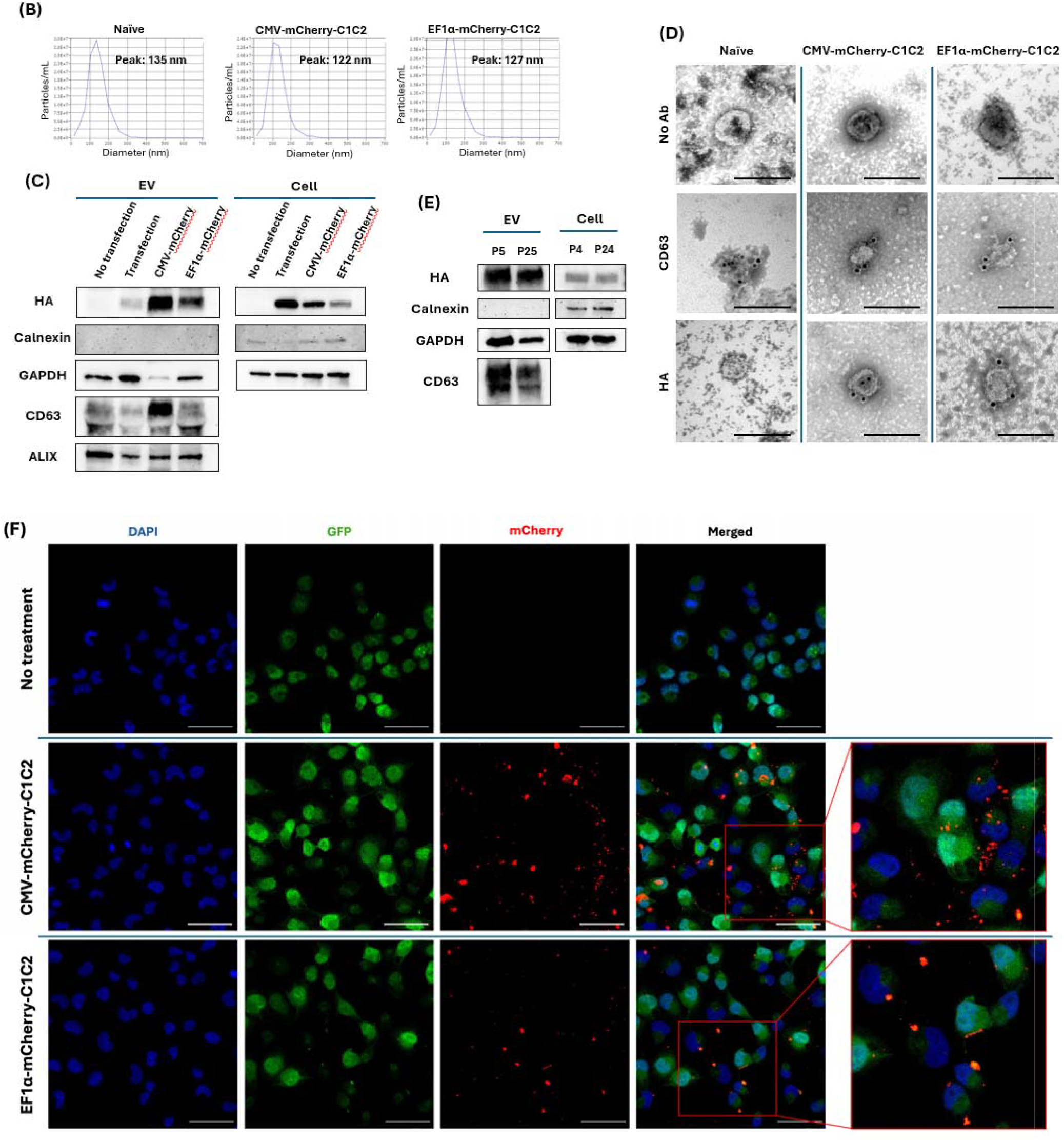
Characterization of the selected stable cell lines and isolated EVs. **(A)** Subcellular localization of mCherry-C1C2 fusion protein in stable cell lines. Representative fluorescence microscopy images of HEK293T cells acquired using Thunder imaging system. Top row: Unmodified HEK293T cells (negative control) showing no mCherry signal. Middle row: CMV-mCherry-C1C2 clone #8. Bottom row: EF-1α-mCherry-C1C2 clone #14. Left column: Brightfield images. Center column: mCherry fluorescence (red channel). Right column: Merged brightfield and fluorescence with higher magnification insets. mCherry-C1C2 displays punctate cytoplasmic localization consistent with endosomal or multivesicular body compartments involved in EV biogenesis. Main images scale bar: 100 μm. **(B)** Nanoparticle Tracking Analysis (NTA) of EV size distribution. Naïve EVs and EVs from CRISPR engineered cells (CMV-mCherry-C1C2 clone #8 and EF-1α-mCherry-C1C2 clone #14). **(C)** Western blot analysis of cellular and EV protein expression with isolated EVs (2×10^9^ particles each: left panel) and cell lysate (40ug each: right panel). Antibodies: anti-HA for mCherry-C1C2 detection, anti-CD63 and anti-ALIX for EV markers, anti-Calnexin for cell contamination assessment, and anti-GAPDH for loading control and PS-binding protein analysis. Note: Gap in CD63 EV blots attributed to albumin co-purification during TFF concentration. **(D)** Single-particle immuno-electron microscopy validation of EV surface engineering. Representative transmission electron microscopy images of EVs from stable cell lines processed for immunogold labeling. Top row: EVs from unmodified HEK293T (negative control), CMV-mCherry-C1C2 clone #8, and EF-1α-mCherry-C1C2 clone #14 incubated with secondary antibody only (no primary antibody control). Middle row: EVs from unmodified HEK293T (negative control), CMV-mCherry-C1C2 clone #8, and EF-1α-mCherry-C1C2 clone #14 labeled with anti-CD63 antibody (EV marker). Bottom row: EV from the same clones labeled with anti-HA antibody detecting surface-displayed mCherry-C1C2. The scale bar represents 200nm. **(E)** Long-term stability analysis of transgene expression. Western blot comparison of mCherry-C1C2 expression in EF-1α-mCherry-C1C2 clone #14 at early (passage 4-5) and late passages (passage 24-25). Both cellular and EV expression levels remain stable over extended culture periods (>25 passages, ~3 months), demonstrating robust transgene maintenance at the AAVS1 locus. GAPDH serves as loading control. **(F)** Functional uptake analysis of engineered EVs. Confocal microscopy images showing uptake of mCherry-labeled EVs (red) by H1299-GFP recipient cells. Top row: CMV-mCherry-C1C2 EVs. Bottom row: EF-1α-mCherry-C1C2 EVs. Cells were incubated with 2×10^8^ EVs for 4 hours at 37°C, washed to remove unbound EVs, and fixed for imaging. DAPI stains nuclei (blue). Colocalization of red and green signals in merged images confirms successful EV internalization. Scale bar: 100 μm. Images acquired using Nikon A1 HD25 confocal microscope at 60× magnification.

EVs isolated from stable cell lines using tangential flow filtration (TFF) maintained characteristic size distributions indistinguishable from naive EVs, with mean diameters ranging from 120-130 nm by nanoparticle tracking analysis (Fig. 3B). This size range is consistent with the expected EV population^3^ and indicates that genetic modification did not alter fundamental EV biophysical properties.

Western blot analysis confirmed the presence of typical EV markers CD63 and ALIX in isolated vesicles (Fig. 3C), consistent with established EV characterization standards^5,6^. The gap observed in CD63 bands was attributed to albumin (components of the EV storage buffer^29^) co-purification during TFF concentration. Importantly, mCherry-C1C2 expression levels in stable cell line-derived EVs exceeded those from transient transfection controls. The inverse correlation between mCherry-C1C2 and GAPDH levels was maintained in the EV fraction, supporting the PS-binding competition mechanism described for GAPDH^42^. Absence of the ER marker calnexin confirmed minimal cellular contamination in EV preparations.

### Single-Particle Validation of EV Surface Engineering

To confirm surface display of the engineered fusion protein, we performed immuno-transmission electron microscopy (immuno-TEM) analysis at the single-particle level (Fig. 3D, Fig. S4). Anti-CD63 labeling confirmed the presence of this EV marker across all vesicle populations. Notably, anti-HA labeling detected surface-displayed mCherry-C1C2 on EVs from stable cell lines but not on control EVs, providing direct evidence of successful surface engineering. The particle sizes observed by electron microscopy correlated well with the size distributions measured by nanoparticle tracking analysis, confirming the morphological integrity of engineered EVs.

### Long-term Stability and Functional Properties

To assess the suitability of stable cell lines for extended production periods, we evaluated transgene expression stability over 25 passages using EF-1α-mCherry-C1C2 clone #14 (Fig. 3E). Both cellular and EV expression levels remained stable over this extended culture period (approximately 3 months), with no appreciable decline in mCherry-C1C2 expression. This stability demonstrates the robust transgene maintenance achieved through AAVS1 integration, consistent with previous reports of stable expression at this locus^36^.

Functional uptake analysis confirmed that engineered EVs retained their natural cellular internalization properties (Fig. 3F). H1299-GFP recipient cells efficiently internalized mCherry-labeled EVs from both CMV and EF-1α promoter-driven cell lines after 4-hour incubation. Colocalization of red (EV) and green (recipient cell) fluorescence signals confirmed successful EV uptake, indicating that surface engineering did not compromise the natural uptake mechanisms of EVs as previously characterized for EV internalization^43,44^.

### Production Consistency and Long-term Stability

The stable expression platform maintained consistent production of uniformly modified EVs from laboratory-scale cultures (2 mL) to larger production volumes (40 mL). Critically, EV populations showed equivalent characteristics across scales, including size distribution (120-130 nm), marker expression (CD63, ALIX), surface modification efficiency, and functional properties (cellular uptake).

This consistency addresses key limitations of transient transfection approaches: (1) heterogeneous EV populations with variable modification efficiency between batches, and (2) loss of transgene expression over time requiring repeated transfections. In contrast, stable cell lines maintained robust mCherry-C1C2 expression over at least 25 passages (~3 months) without decline, enabling continuous production of uniformly modified EVs from a single engineered cell line. No significant differences in EV characteristics were observed between constructs driven by CMV versus EF-1α promoters, though the EF-1α system showed higher overall integration efficiency and more diverse clone populations including homozygous integrations. These results establish a robust, reproducible platform for generating stable cell lines capable of sustained, consistent production of surface-modified EVs with reduced batch-to-batch variability suitable for applications requiring uniform EV populations over extended manufacturing periods.

## Discussion

This study establishes a CRISPR-Cas9-based method for generating stable cell lines that produce surface-engineered EVs through targeted integration of mCherry-C1C2 constructs into the AAVS1 locus. Our approach provides a reproducible platform for consistent EV production that addresses some of the variability issues associated with transient transfection methods^22,26^. The data demonstrated that stable cell lines exhibited enhanced surface modification efficiency relative to transient transfection, despite reduced intracellular expression levels. This unexpected outcome indicated intrinsic differences in trafficking and sorting pathways between constitutive and transient expression systems. One possible explanation for the enhanced EV incorporation efficiency in stable cell lines is that transient expression, while generating higher total protein levels, may overwhelm cellular trafficking machinery, leading to intracellular accumulation rather than efficient secretion into EVs. Interestingly, the inverse correlation between mCherry-C1C2 and GAPDH incorporation into EVs provides mechanistic insight into phosphatidylserine-mediated competition for surface display, suggesting that stable expression enhances the efficiency of EV loading by optimizing cellular machinery rather than by elevating protein abundance.

A critical advantage of the stable cell line platform is the sustained production capability that eliminates the need for repeated transfections. Transient transfection systems lose transgene expression within days to weeks, requiring continuous re-transfection cycles that introduce batch-to-batch variability and labor-intensive workflows. In contrast, our stable cell lines maintained robust mCherry-C1C2 expression over at least 25 passages (~3 months) without decline (Fig. 3E), enabling continuous production from a single engineered line. Additionally, while transient methods produce heterogeneous EV populations where only ~50% of EVs display the intended modification^23^, the stable expression platform reduces this heterogeneity, producing more uniform EV populations essential for therapeutic applications requiring reproducible product specifications. The combination of long-term expression stability and consistent modification efficiency positions this platform as a manufacturing solution that addresses both the temporal and qualitative challenges of EV therapeutic production.

In stable cell lines, the preferential localization of mCherry-C1C2 to endosomal and multivesicular body compartments likely accounts for the enhanced efficiency of EV surface modification. The C1C2 domain’s high affinity for phosphatidylserine appears to drive efficient trafficking toward EV-forming compartments, particularly under constitutive expression conditions where cellular sorting machinery can optimize protein distribution. This contrasts with alternative approaches such as Lamp2B fusion proteins, which are prone to lysosomal sorting and N-terminal peptide instability^13,45^. The PS-binding strategy thus provides a mechanistic advantage for EV surface engineering by leveraging the natural phosphatidylserine externalization on EV membranes.

The competition between mCherry-C1C2 and endogenous GAPDH for PS-binding sites reveals mechanistic aspects of EV surface modification^42^. GAPDH is known to bind PS and localize on EV surfaces^42,46^, positioning it as a natural competitor for engineered PS-binding proteins. Our data suggest that in stable cell lines, constitutive expression of mCherry-C1C2 can displace GAPDH from PS-binding sites, potentially reducing heterogeneity and promoting more uniform surface modification across EV populations. Stable expression may optimize EV loading through altered protein competition dynamics rather than simply increasing protein availability. The distinct trafficking routes may contribute to this effect: LAMP2B primarily undergoes lysosomal sorting^13^, whereas the PS-binding affinity of the C1C2 domain appears to favor trafficking toward EV-generating compartments^22,47^.

The CRISPR-Cas9 integration process followed established protocols, with puromycin selection effectively enriching for cells containing the resistance cassette. Following FACS sorting of mCherry-positive cells into single-cell clones, genotyping PCR confirmed successful gene insertion in 7 out of 17 CMV-driven clones and 35 out of 45 EF-1α-driven clones (combining 12 heterozygous and 23 homozygous insertions). The genotyping strategy using junction PCR and insertion site PCR effectively distinguished between heterozygous and homozygous integration events, consistent with established AAVS1 targeting protocols^35,41^. Both promoter systems supported stable expression over at least 25 passages, with no apparent growth disadvantages compared to parental cells, confirming the utility of AAVS1 as a genomic safe harbor^35,36^. EVs produced from stable cell lines maintained typical characteristics including size distribution (120-130 nm), marker expression (CD63, ALIX), and functional properties (cellular uptake)^43,44^. The method successfully generated EVs with consistent surface modification, as confirmed by immuno-TEM single-particle analysis. The platform demonstrated scalability from laboratory volumes to larger culture systems with maintained product quality.

This method offers several advantages over transient transfection approaches: (1) elimination of repeated transfection procedures, (2) reduced batch-to-batch variability, (3) stable long-term production capability, and (4) potential for industrial scaling. The use of AAVS1 as the integration site minimizes concerns about positional effects that can compromise expression stability in random integration systems^32,33^. Additionally, the stable cell line platform addresses storage-related reproducibility issues that we previously identified as significant barriers to EV therapeutic development^29^. However, several limitations warrant consideration. While surface modifications were confirmed at the single particle level, quantification of modification density per unit EV surface area remains challenging and would require more sensitive analytical techniques. The relationship between cellular expression levels and EV incorporation efficiency requires further investigation to optimize production parameters for different applications.

Chemical modification approaches, such as click chemistry^24^ or lipid-anchored ligand insertion^48^, can achieve consistent modifications but face limitations in scalability and may alter EV biophysical properties^48^. Our genetic approach using stable cell lines combines the specificity of genetic engineering with improved consistency compared to transient transfection systems^23^. The observed reduction in modification heterogeneity suggests this platform could produce more uniform products suitable for therapeutic applications^27,28^. The choice of AAVS1 as the integration site proved advantageous, as evidenced by stable expression without apparent cellular toxicity or growth disadvantages^32,49^. This contrasts with random integration approaches that suffer from positional effects and epigenetic silencing^32,36^. The robust expression observed across multiple clones suggests that AAVS1-targeted integration provides a reliable foundation for industrial EV production.

Approaches to increase EV production include the use of bioreactors such as hollow fiber or hyperflask systems, as well as methods to stimulate cells and enhance EV secretion^50-53^. The stable expression platform developed here could be readily adapted to such production-scale systems. Combined with efficient purification methods such as tangential flow filtration followed by size exclusion chromatography (TFF-SEC)^54^, this approach provides a foundation for consistent large-scale EV production. The platform’s versatility allows for adaptation to different applications by substituting alternative targeting ligands or incorporating different functional domains. This modularity could facilitate the development of specialized EV products while maintaining the consistency advantages of stable expression^11,27^.

The current study focused on proof-of-concept using mCherry-C1C2 as a model system; adaptation to therapeutic applications will require systematic evaluation of different targeting ligands and cargo loading strategies. Understanding the precise mechanisms governing PS-mediated protein sorting into EVs could enable rational optimization of engineering efficiency^35,42^.

Future applications may include the incorporation of therapeutic proteins, nucleic acids, or targeting ligands for specific cell types or disease states^11,55^. The stable expression platform could facilitate the development of personalized EV therapeutics by enabling consistent production from patient-derived cells. Additionally, the method could be extended to other cell types beyond HEK293T to leverage cell-specific EV properties for particular applications^10^.

## Conclusion

By inserting different targeting molecules into the AAVS1 locus, this method enables stable production of engineered EVs tailored to specific applications. The stable cell line platform provides efficient EV surface modification and represents a scalable technology for consistent production. Combined with efficient purification methods^54^, this approach offers a foundation for EV-based applications requiring reproducible and uniform surface-modified vesicles.

## Acknowledgement

We thank Dr. Jens Schmidt for valuable guidance on this work, the Center for Advanced Microscopy for confocal imaging, and the IQ Flowcytometry Core for NTA at Michigan State University. Graphical abstract was created with BioRender.com. TS is supported by the Eunice Kennedy Shriver National Institute of Child Health & Human Development of the National Institutes of Health under Award Number T32HD087166. This work was funded in part by 1R21GM154180-01 and 1R01CA286786-01 (MH).

## References

1 Abels, E. R. B. X. O. Introduction to Extracellular Vesicles: Biogenesis, RNA Cargo Selection, Content, Release, and Uptake. (Cell Mol Neurobio, 2016).

2 Fuhrmann, G., Herrmann, I. K. & Stevens, M. M. Cell-derived vesicles for drug therapy and diagnostics: opportunities and challenges. Nano Today 10, 397–409, doi:10.1016/j.nantod.2015.04.004 (2015).

3 Doyle, L. M. & Wang, M. Z. Overview of Extracellular Vesicles, Their Origin, Composition, Purpose, and Methods for Exosome Isolation and Analysis. Cells 8, doi:10.3390/cells8070727 (2019).

4 Colombo, M. M., C. van Niel, G. Kowal, J. Vigneron, J. Benaroch, P. Manel, N. Moita, L. F. Théry, C. Raposo, G. Analysis of ESCRT functions in exosome biogenesis, composition and secretion highlights the heterogeneity of extracellular vesicles. (J Cell Sci, 2013).

5 Kowal, J. et al. Proteomic comparison defines novel markers to characterize heterogeneous populations of extracellular vesicle subtypes. Proc Natl Acad Sci U S A 113, E968–977, doi:10.1073/pnas.1521230113 (2016).

6 Sinha, A., Ignatchenko, V., Ignatchenko, A., Mejia-Guerrero, S. & Kislinger, T. In-depth proteomic analyses of ovarian cancer cell line exosomes reveals differential enrichment of functional categories compared to the NCI 60 proteome. Biochem Biophys Res Commun 445, 694–701, doi:10.1016/j.bbrc.2013.12.070 (2014).

7 Hallal, S., Tűzesi, Á., Grau, G. E., Buckland, M. E. & Alexander, K. L. Understanding the extracellular vesicle surface for clinical molecular biology. J Extracell Vesicles 11, e12260, doi:10.1002/jev2.12260 (2022).

8 Matsumura, S. et al. Subtypes of tumour cell-derived small extracellular vesicles having differently externalized phosphatidylserine. J Extracell Vesicles 8, 1579541, doi:10.1080/20013078.2019.1579541 (2019).

9 Miyanishi, M. et al. Identification of Tim4 as a phosphatidylserine receptor. Nature 450, 435–439, doi:10.1038/nature06307 (2007).

10 Maumus, M., Rozier, P., Boulestreau, J., Jorgensen, C. & Noel, D. Mesenchymal Stem Cell-Derived Extracellular Vesicles: Opportunities and Challenges for Clinical Translation. Front Bioeng Biotechnol 8, 997, doi:10.3389/fbioe.2020.00997 (2020).

11 Komuro, H., Aminova, S., Lauro, K. & Harada, M. Advances of engineered extracellular vesicles-based therapeutics strategy. Sci Technol Adv Mater 23, 655–681, doi:10.1080/14686996.2022.2133342 (2022).

12 Tang, F. D., Tao Zhou, Chengqian Deng, Leon Liu, Hans B. Wang, Wenshen Liu, Guanshu Ying, Mingyao Li, Pan P. Genetically engineered human induced pluripotent stem cells for the production of brain-targeting extracellular vesicles. (Springer Science and Business Media LLC, 2024).

13 Li, Z. et al. Fusion protein engineered exosomes for targeted degradation of specific RNAs in lysosomes: a proof-of-concept study. J Extracell Vesicles 9, 1816710, doi:10.1080/20013078.2020.1816710 (2020).

14 Alvarez-Erviti, L. et al. Delivery of siRNA to the mouse brain by systemic injection of targeted exosomes. Nat Biotechnol 29, 341–345, doi:10.1038/nbt.1807 (2011).

15 Tian, Y. et al. A doxorubicin delivery platform using engineered natural membrane vesicle exosomes for targeted tumor therapy. Biomaterials 35, 2383–2390, doi:10.1016/j.biomaterials.2013.11.083 (2014).

16 Wang, X. et al. Engineered Exosomes With Ischemic Myocardium-Targeting Peptide for Targeted Therapy in Myocardial Infarction. J Am Heart Assoc 7, e008737, doi:10.1161/JAHA.118.008737 (2018).

17 Scott, T. A. et al. Engineered extracellular vesicles directed to the spike protein inhibit SARS-CoV-2. Mol Ther Methods Clin Dev 24, 355–366, doi:10.1016/j.omtm.2022.01.015 (2022).

18 Yao, X. et al. Engineered extracellular vesicles as versatile ribonucleoprotein delivery vehicles for efficient and safe CRISPR genome editing. J Extracell Vesicles 10, e12076, doi:10.1002/jev2.12076 (2021).

19 Cone, A. S. et al. CD81 fusion alters SARS-CoV-2 Spike trafficking. mBio 15, e0192224, doi:10.1128/mbio.01922-24 (2024).

20 Mai, J., Wang, K., Liu, C., Xiong, S. & Xie, Q. αvβ3-targeted sEVs for efficient intracellular delivery of proteins using MFG-E8. BMC Biotechnol 22, 15, doi:10.1186/s12896-022-00745-7 (2022).

21 Delcayre, A. et al. Exosome Display technology: applications to the development of new diagnostics and therapeutics. Blood Cells Mol Dis 35, 158–168, doi:10.1016/j.bcmd.2005.07.003 (2005).

22 Komuro, H. et al. Engineering Extracellular Vesicles to Target Pancreatic Tissue In Vivo. Nanotheranostics 5, 378–390, doi:10.7150/ntno.54879 (2021).

23 Silva, A. M. et al. Quantification of protein cargo loading into engineered extracellular vesicles at single-vesicle and single-molecule resolution. J Extracell Vesicles 10, e12130, doi:10.1002/jev2.12130 (2021).

24 Smyth, T. et al. Surface functionalization of exosomes using click chemistry. Bioconjug Chem 25, 1777–1784, doi:10.1021/bc500291r (2014).

25 Liu, Q., Li, D., Pan, X. & Liang, Y. Targeted therapy using engineered extracellular vesicles: principles and strategies for membrane modification. J Nanobiotechnology 21, 334, doi:10.1186/s12951-023-02081-0 (2023).

26 Komuro, H., Aminova, S., Lauro, K., Woldring, D. & Harada, M. Design and Evaluation of Engineered Extracellular Vesicle (EV)-Based Targeting for EGFR-Overexpressing Tumor Cells Using Monobody Display. Bioengineering (Basel) 9, doi:10.3390/bioengineering9020056 (2022).

27 Chang, R. K., Mathias, N. & Hussain, M. A. Biopharmaceutical Evaluation and CMC Aspects of Oral Modified Release Formulations. AAPS J 19, 1348–1358, doi:10.1208/s12248-017-0112-6 (2017).

28 Witwer, K. W. & Wolfram, J. Extracellular vesicles versus synthetic nanoparticles for drug delivery. Nat Rev Mater 6, 103–106, doi:10.1038/s41578-020-00277-6 (2021).

29 Kawai-Harada, Y., El Itawi, H., Komuro, H. & Harada, M. Evaluation of EV Storage Buffer for Efficient Preservation of Engineered Extracellular Vesicles. Int J Mol Sci 24, doi:10.3390/ijms241612841 (2023).

30 Takakura, Y. et al. Quality and Safety Considerations for Therapeutic Products Based on Extracellular Vesicles. Pharm Res 41, 1573–1594, doi:10.1007/s11095-024-03757-4 (2024).

31 Verma, N. & Arora, S. Navigating the Global Regulatory Landscape for Exosome-Based Therapeutics: Challenges, Strategies, and Future Directions. Pharmaceutics 17, doi:10.3390/pharmaceutics17080990 (2025).

32 Smith, J. R. et al. Robust, persistent transgene expression in human embryonic stem cells is achieved with AAVS1-targeted integration. Stem Cells 26, 496–504, doi:10.1634/stemcells.2007-0039 (2008).

33 Cabrera, A. et al. The sound of silence: Transgene silencing in mammalian cell engineering. Cell Syst 13, 950–973, doi:10.1016/j.cels.2022.11.005 (2022).

34 Dubois, V. P. et al. Safe Harbor Targeted CRISPR-Cas9 Tools for Molecular-Genetic Imaging of Cells in Living Subjects. CRISPR J 1, 440–449, doi:10.1089/crispr.2018.0030 (2018).

35 Shin, S. et al. Comprehensive Analysis of Genomic Safe Harbors as Target Sites for Stable Expression of the Heterologous Gene in HEK293 Cells. ACS Synth Biol 9, 1263–1269, doi:10.1021/acssynbio.0c00097 (2020).

36 Oceguera-Yanez, F. et al. Engineering the AAVS1 locus for consistent and scalable transgene expression in human iPSCs and their differentiated derivatives. Methods 101, 43–55, doi:10.1016/j.ymeth.2015.12.012 (2016).

37 Natsume, T., Kiyomitsu, T., Saga, Y. & Kanemaki, M. T. Rapid Protein Depletion in Human Cells by Auxin-Inducible Degron Tagging with Short Homology Donors. Cell Rep 15, 210–218, doi:10.1016/j.celrep.2016.03.001 (2016).

38 Broadbent, D. G., Barnaba, C., Perez, G. I. & Schmidt, J. C. Quantitative analysis of autophagy reveals the role of ATG9 and ATG2 in autophagosome formation. J Cell Biol 222, doi:10.1083/jcb.202210078 (2023).

39 Liu, X. et al. The influence of polymeric properties on chitosan/siRNA nanoparticle formulation and gene silencing. Biomaterials 28, 1280–1288, doi:10.1016/j.biomaterials.2006.11.004 (2007).

40 Alamuru-Yellapragada, N. P., Kapadia, B. & Parsa, K. V. L. In-house made nucleofection buffer for efficient and cost effective transfection of RAW 264.7 macrophages. Biochem Biophys Res Commun 487, 247–254, doi:10.1016/j.bbrc.2017.04.043 (2017).

41 Qian, K. et al. A simple and efficient system for regulating gene expression in human pluripotent stem cells and derivatives. Stem Cells 32, 1230–1238, doi:10.1002/stem.1653 (2014).

42 Dar, G. H. et al. GAPDH controls extracellular vesicle biogenesis and enhances the therapeutic potential of EV mediated siRNA delivery to the brain. Nat Commun 12, 6666, doi:10.1038/s41467-021-27056-3 (2021).

43 Bonsergent, E. et al. Quantitative characterization of extracellular vesicle uptake and content delivery within mammalian cells. Nat Commun 12, 1864, doi:10.1038/s41467-021-22126-y (2021).

44 Toribio, V. et al. Development of a quantitative method to measure EV uptake. Sci Rep 9, 10522, doi:10.1038/s41598-019-47023-9 (2019).

45 Hung, M. E. & Leonard, J. N. Stabilization of exosome-targeting peptides via engineered glycosylation. J Biol Chem 290, 8166–8172, doi:10.1074/jbc.M114.621383 (2015).

46 Kaneda, M., Takeuchi, K., Inoue, K. & Umeda, M. Localization of the phosphatidylserine-binding site of glyceraldehyde-3-phosphate dehydrogenase responsible for membrane fusion. J Biochem 122, 1233–1240, doi:10.1093/oxfordjournals.jbchem.a021886 (1997).

47 Shi, J., Heegaard, C. W., Rasmussen, J. T. & Gilbert, G. E. Lactadherin binds selectively to membranes containing phosphatidyl-L-serine and increased curvature. Biochim Biophys Acta 1667, 82–90, doi:10.1016/j.bbamem.2004.09.006 (2004).

48 Liu, Y. et al. Engineered Extracellular Vesicles for Delivery of an IL-1 Receptor Antagonist Promote Targeted Repair of Retinal Degeneration. Small 19, doi:10.1002/smll.202302962 (2023).

49 Luo, Y. et al. Stable enhanced green fluorescent protein expression after differentiation and transplantation of reporter human induced pluripotent stem cells generated by AAVS1 transcription activator-like effector nucleases. Stem Cells Transl Med 3, 821–835, doi:10.5966/sctm.2013-0212 (2014).

50 Grangier, A. et al. Technological advances towards extracellular vesicles mass production. Adv Drug Deliv Rev 176, 113843, doi:10.1016/j.addr.2021.113843 (2021).

51 Mendt, M. et al. Generation and testing of clinical-grade exosomes for pancreatic cancer. JCI Insight 3, doi:10.1172/jci.insight.99263 (2018).

52 Parolini, I. et al. Microenvironmental pH is a key factor for exosome traffic in tumor cells. J Biol Chem 284, 34211–34222, doi:10.1074/jbc.M109.041152 (2009).

53 Yan, L. & Wu, X. Exosomes produced from 3D cultures of umbilical cord mesenchymal stem cells in a hollow-fiber bioreactor show improved osteochondral regeneration activity. Cell Biol Toxicol 36, 165–178, doi:10.1007/s10565-019-09504-5 (2020).

54 Kawai-Harada, Y., Nimmagadda, V. & Harada, M. Scalable isolation of surface-engineered extracellular vesicles and separation of free proteins via tangential flow filtration and size exclusion chromatography (TFF-SEC). BMC Methods 1, 9, doi:10.1186/s44330-024-00009-0 (2024).

55 Tang, F. et al. Genetically engineered human induced pluripotent stem cells for the production of brain-targeting extracellular vesicles. Stem Cell Research & Therapy 15, doi:10.1186/s13287-024-03955-2 (2024).

