## Supplemental figures for "Generation of Cellular Biofactories for Scalable Production of Surface-Engineered Extracellular Vesicles via CRISPR Genome Editing"

Supplemental Information

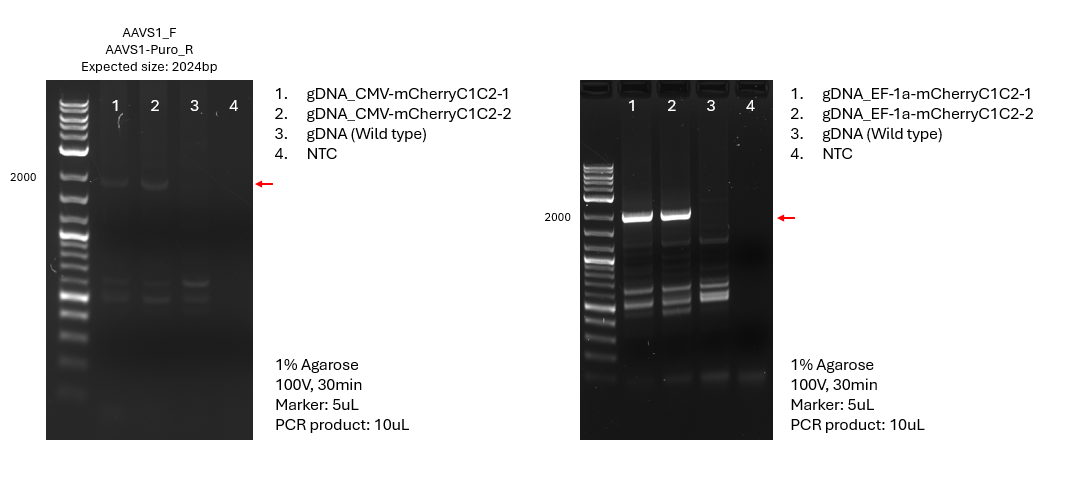

**Figure S1. Validation of CRISPR-mediated gene integration in bulk cell populations**. PCR analysis confirming successful CRISPR-Cas9-mediated integration of mCherry-C1C2 constructs into bulk cell populations following puromycin selection. Genomic DNA was extracted from puromycin-selected cell pools and analyzed using primers spanning the junction site between the integrated construct and the AAVS1 locus. The presence of the expected PCR product (red arrow) confirms enrichment of CRISPR-positive cells in the bulk population prior to single-cell sorting. Lane assignments and molecular weight markers are indicated.

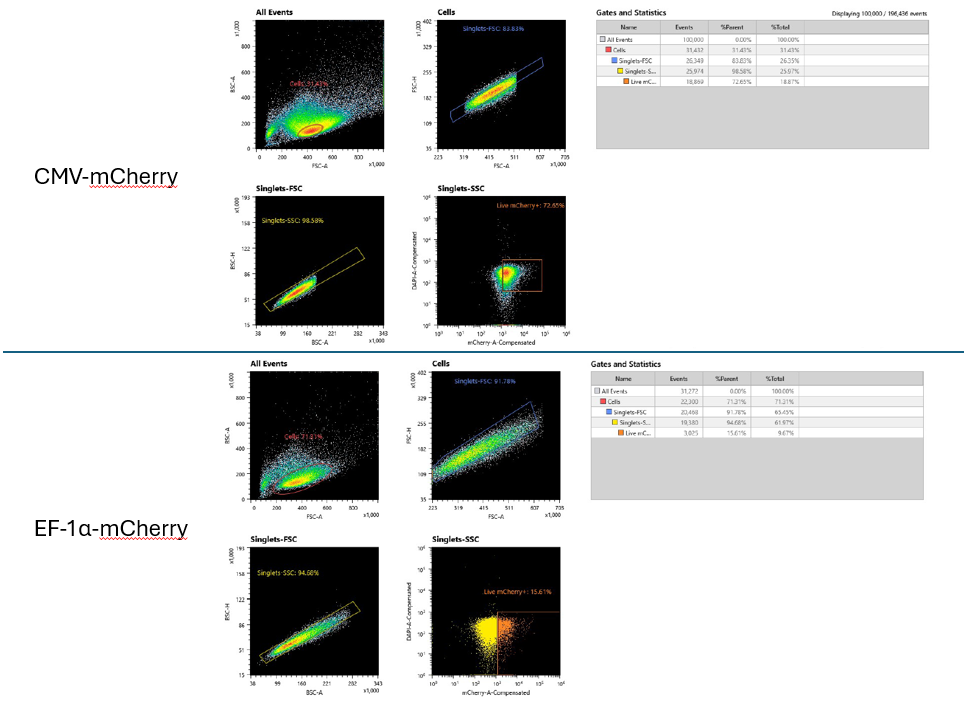

**Figure S2. Fluorescence-activated cell sorting strategy for single-cell isolation**. FACS analysis and gating strategy used to isolate single mCherry-positive cells from CRISPR-edited bulk populations. Cells were stained with DAPI to exclude dead cells, and viable cells (DAPI-negative) expressing mCherry fluorescence were identified and single-cell sorted into 96-well plates. The gating strategy shows the sequential selection process: (top-left) forward scatter area vs. side scatter area for cell population identification, (top-right) forward scatter area vs. forward scatter height for singlets of cell population identification, (bottom-left) side scatter area vs. side scatter height for singlets of cell population identification, and (bottom-right) DAPI exclusion for viability, and mCherry-positive cell selection for single-cell sorting.

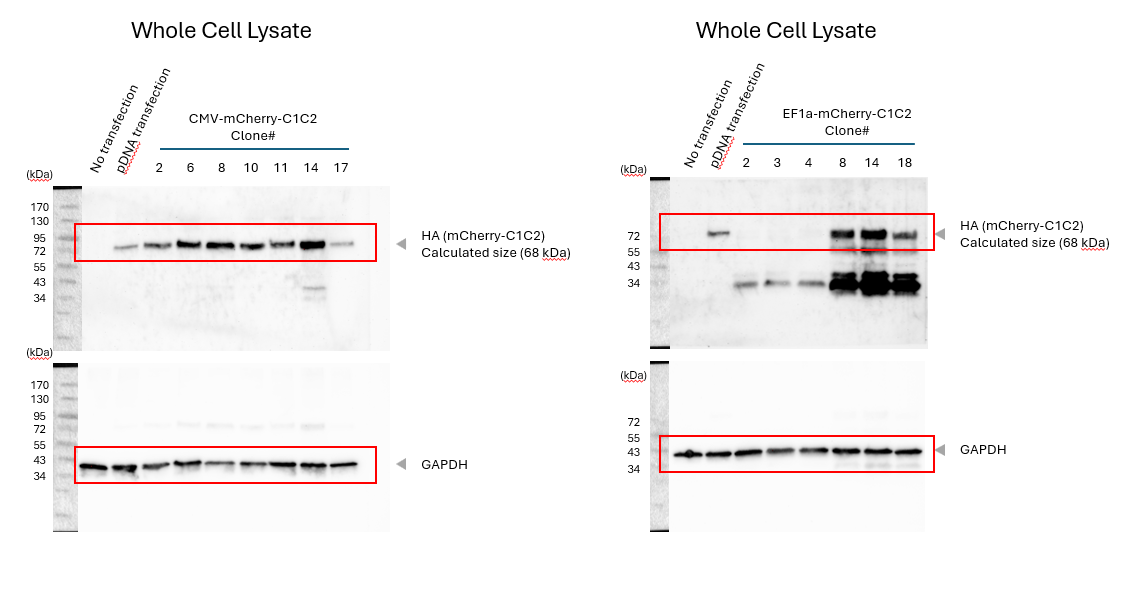

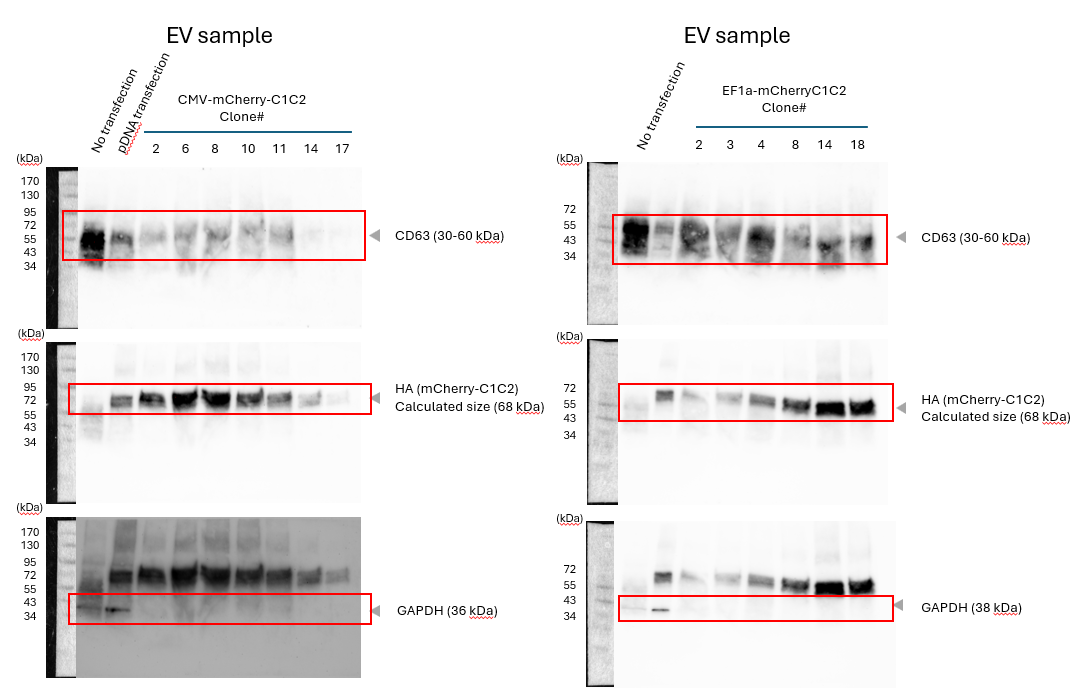

**Figure S3. Western blot analysis of clone characterization.** Full western blot images corresponding to Figure 2B, showing mCherryC1C2 expression analysis in selected clones. Top panel: Cell lysate (40 µg protein per lane). Bottom panel: EVs (equivalent to 15 µg protein per lane). The EF-1α promoter-driven constructs show expression of N-terminal fragments of the mCherry-C1C2 fusion protein (indicated by red arrow), likely resulting from alternative translation initiation or post-translational processing. These fragments do not affect EV surface engineering efficiency. Red squares indicate the cropped regions used in the main Figure 2B. Antibodies used: anti-HA for mCherry-C1C2 detection, anti-GAPDH for loading control and PS-binding protein analysis.

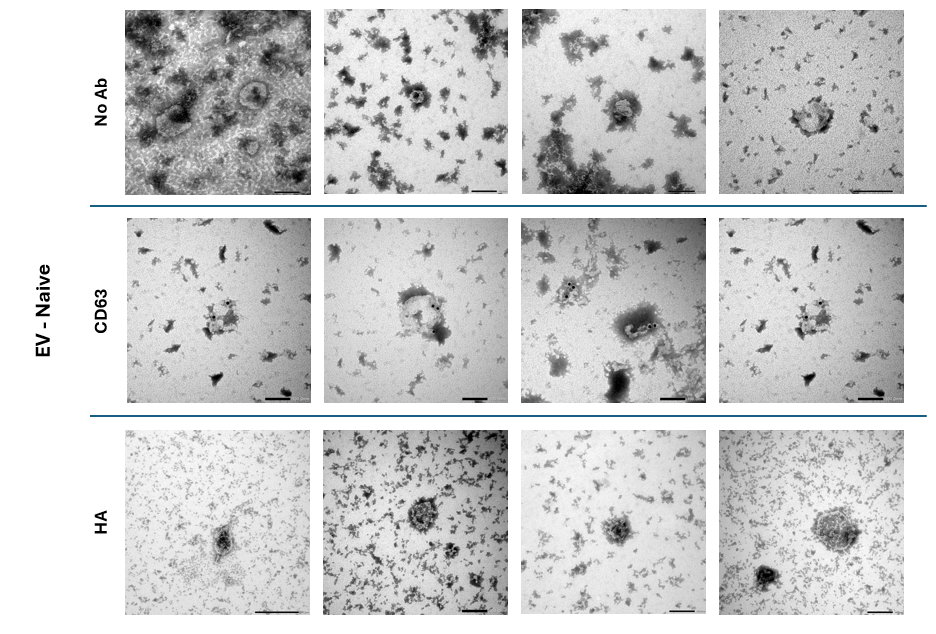

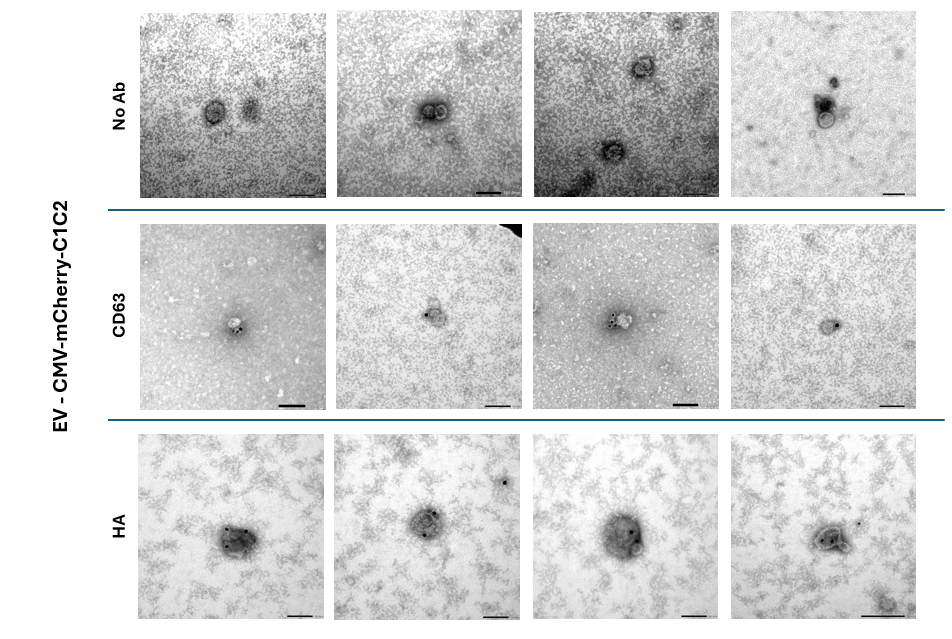

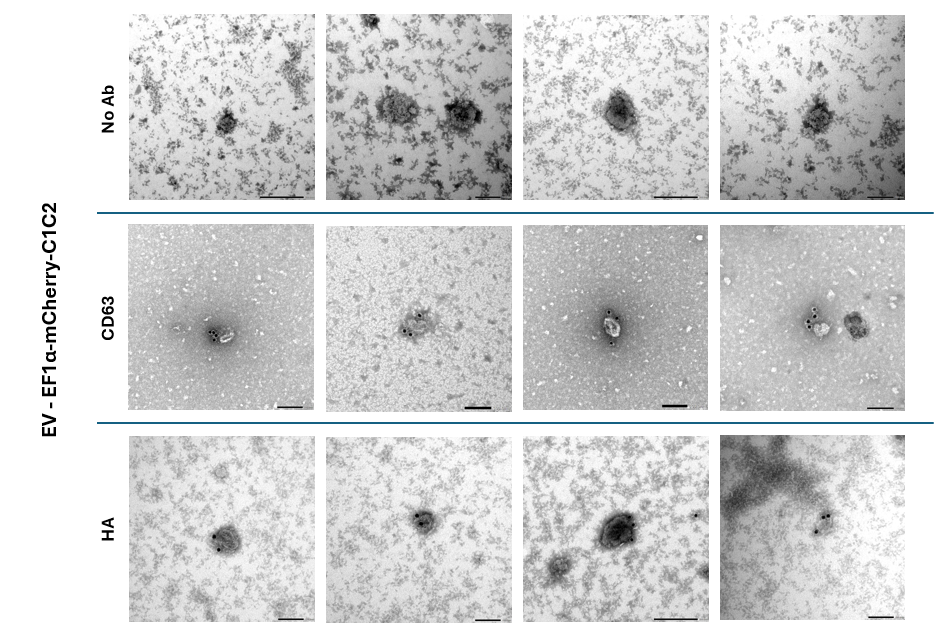

**Figure S4. Single-particle immuno-electron microscopy validation.** Extended collection of transmission electron microscopy images showing immunogold labeling of EVs from stable cell lines. EVs were processed for immunogold labeling using anti-CD63 (EV marker) and anti-HA (detecting surface-displayed mCherry-C1C2) antibodies. Images demonstrate consistent surface modification across multiple individual vesicles from both CMV-mCherry-C1C2 and EF-1α-mCherry-C1C2 stable cell lines. The presence of gold particles on EV surfaces confirms successful display of the engineered fusion protein. Control EVs from unmodified HEK293T cells show CD63 labeling but absence of HA labeling, validating the specificity of surface engineering. Scale bars represent 200 nm.

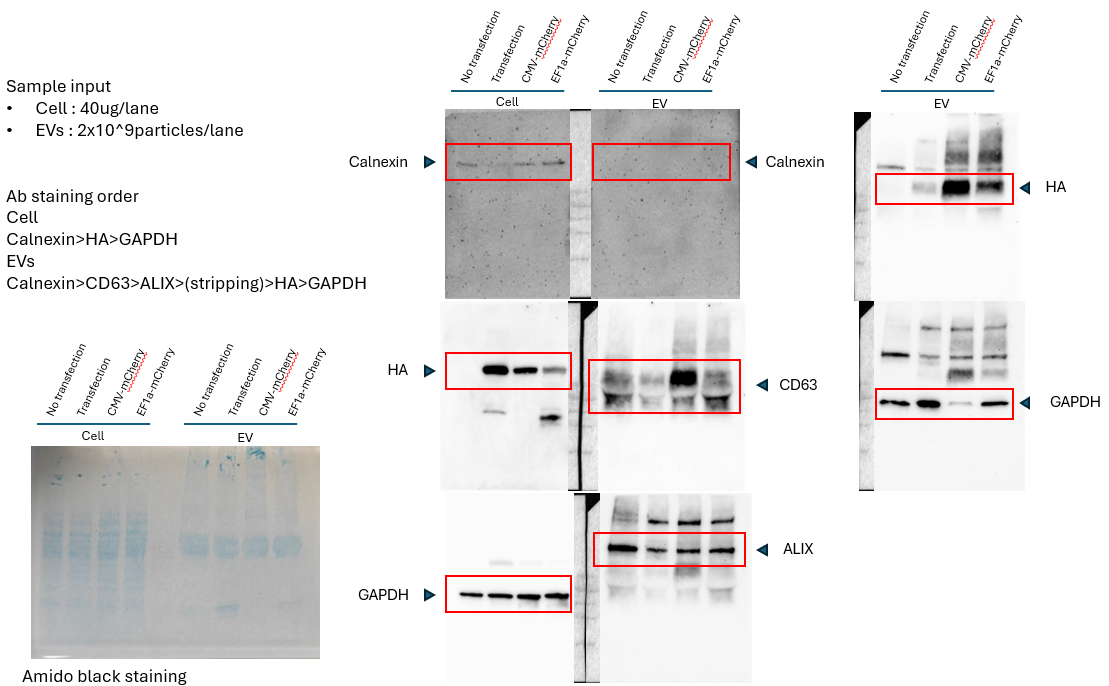

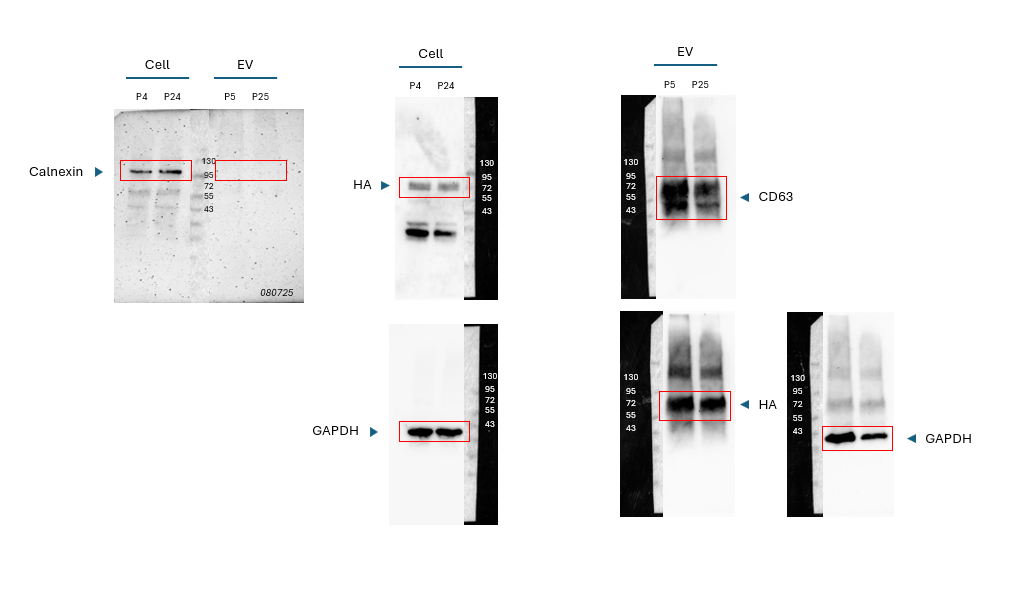

Figure S5. **Complete western blot images for EV characterization and stability analysis.** Full Western blot images corresponding to Figure 3C (EV characterization) and Figure 3E (long-term stability analysis). Top panels: Complete blots showing EV markers (CD63, ALIX), cellular contamination marker (calnexin), and engineered protein (mCherry-C1C2 detected by anti-HA) in EVs isolated from stable cell lines and transient transfection controls. Bottom panels: Long-term expression stability analysis showing maintained mCherry-C1C2 expression levels in both cells and EVs after extended passaging (passage 4-5 vs. passages 24-25) of EF-1α-mCherry-C1C2 clone #14. GAPDH serves as loading control for cellular lysates and as an endogenous PS-binding protein control for EVs. The gap in CD63 EV blots is attributed to albumin co-purification during tangential flow filtration concentration.
